# In-vitro study of the activity of some medicinal plant leaf extracts on urinary tract infection causing bacterial pathogens isolated from indigenous people of Bolangir district, Odisha, India

**DOI:** 10.1101/2020.06.25.172650

**Authors:** Mirkashim H. Saha, Saumya Dash, Abhisek Mishra, Jyotismita Satpathy, Sujit K. Mishra

**Affiliations:** Laboratory of Natural products & Therapeutics, School of Applied Sciences, Centurion University of Technology and Management, Odisha, India; Stem Cell Facility, All India Institute of Medical Sciences, New Delhi, India; Nabakrushna Choudhury Centre for Development Studies, Odisha, India

**Keywords:** Urinary tract infection, Antibiotic resistance, Bacterial pathogens of UTI, Alternative medicines, Plant extract, Therapeutics

## Abstract

Thousands of people over the world are suffering from Urinary tract infection (UTI) each day. Side effects associated with the most preferred allopathic treatment for UTI and antibiotic resistance shown by the pathogens has evolved as major challenge. Owing to this, the present study was carried out to identify the bacterial UTI pathogens among indigenous people of Bolangir district, Odisha, India *vis-à-vis* explore the alternative source of natural therapeutics. Bacterial pathogens causing the UTIs were identified using the colony morphology, Gram’s staining and biochemical characterization techniques. Microbial inhibitory test of antibiotics and leaf extracts were performed using the disc diffusion and well diffusion techniques respectively. MIC and MLC were determined by broth dilution method. Binary probit model was used to determine the prevalence of UTI across people of different age group and gender. Four bacterial strains namely *Escherichia coli*, *Enterococcus faecalis*, *Staphylococcus aureus*, and *Klebsiella pneumoniae* were identified as the causative agents of UTI among the people. The *E. coli* was identified as most infectious while *S. aureus* as the least infectious pathogen. Females in the age group of ‘16-30 years’ and male within ‘61-75 years’ were more susceptible to UTI. Among the tested leaf extracts, *Tamarindus indica* and *Clitoria ternatea* were more effective to treat UTI as compared to the tested antibiotics Ceftriaxone and Piperacillin. The leaf extracts of *T. indica* and *C. ternatea* may effectively be utilized for herbal drug development for treatment of UTI.

## INTRODUCTION

Urinary tract infections (UTIs) are the common bacterial infections usually affects all parts of the urinary system. It is the second most widespread infection after respiratory tract infection (1) affecting 150 million people every year with a higher susceptibility rate of female to this infection than male (2). This occurs most frequently between the age group of 16 and 35 years (3). The infection occurs when bacteria enters the urethra. Infections either limited to the urethra (urethritis) or travel up to bladder and multiply there and cause bladder infection (cystitis). If the infection cannot be treated immediately then the bacteria may travels further upward to the urethers, multiply their and infect the kidneys (pyelonephritis). Most of the common bacterial pathogens of UTI include *Escherichia coli, Klebsiella pneumoniae*, *Staphyloccous aureus*, *Enterobacter* spp., and *Pseudomonas aeruginosa* (2) and about 80% of the UTIs are caused by *E. coli* bacteria alone (1).

Due to its immediate effect, allopathic treatment is mostly preferred to treat this infectious disease. In the allopathic treatments, oral antibiotics like sulfamethoxazole, nitrofurantoin, fosfomycin, amoxicillin and levofloxacin are mostly used (4). However, some side effects such as nausea, diarrhea, dizziness, lightheadedness, headache and/or trouble sleeping are usually encountered in a number of patients while using these antibiotics. Additionally, antibiotic resistance shown by the pathogens has evolved as an emerging problem and has been recognized as a major challenge for the medical practitioner and researchers to prevent these pathogens from causing infection. Therefore, it is imperative to search for new antimicrobials to prevent this infection. The production of synthetic drugs is more expensive and they shows adverse side effects as compared to plant derived natural drugs (5). The plant derived antimicrobial drugs are of natural origin, and have either less or no adverse side effects. Therefore, the natural drugs have a growing demand in the current scenario and could be used as biological control agent against the pathogens.

In Odisha, UTI is prevalent in both male and female of different age groups (2). Kontiokari et al. (6) in a study on women finds that dietary factors are associated with the prevalence UTI. The Odisha state of India in general and Kalahandi-Bolangir-Koraput zone in particular are popularized as the region with the characteristics of poor nutritional intake (7). Therefore, it is expected that the people of this region have a greater exposure to UTI and requires more attention to redress the above health issue. This provides the motivation to examine the UTI status in Bolangir district of Odisha, India *vis-a-vis* explore a new dimension of development of plant derived anti microbial therapeutics as an alternative approach against side effects of allopathic treatment.

The present study was carried out to identify the bacterial pathogens causing urinary tract infection (UTI) *vis-à-vis* determine the prevalence of UTI and the effect of gender and age on its prevalence. This study further evaluated the leaf extracts of 10 medicinal plants of ethano-medicinal importance to determine their effects against the isolated bacterial pathogens causing UTIs.

## MATERIALS AND METHODOLOGY

### Study Population

A total of 602 numbers of patients (both male and female) registered during the period from 12^th^ January 2019 to 30^th^ October 2019 in different wards of hospitals, outdoor patients department, and health care centers in Bolangir district of Odisha, India constitutes the population of the study.

### Collection of urine samples

The urine samples were collected from patients of all ages and sexes. The patients were instructed to collect medium-jet urine after cleaning the genital area. The first jets were discarded and the medium jets were collected into appropriate sterile vials. The samples collected were transported in thermal boxes under refrigerated condition at 4°C and processed immediately after received at the laboratory for characterization.

### Colony count

The urine samples were homogenized and seeding was carried out using inoculation loop (1μl) without any centrifugation. The loops were first vertically deeped in the urine samples and seeded for quantification onto petri-dishes containing cysteine lactose electrolyte-deficient agar medium (8) and incubated at 37°C for 24 hours. The numbers of colonies formed were counted as colony forming unit (CFU) per ml of urine.

### Inclusion criteria

The present study follows Hooton and Stamm (9) and includes male and female of all ages, whose urine culture was positive for UTI and a colony count ≥10^3^ CFU/ml. Based on this criteria, out of 602 patients 162 were excluded. Thus, the study was based on a sample of 440 (male=286, Female=154) UTI registered patients.

### Isolation of the pathogenic bacteria

Isolation of the pathogenic bacteria from the 440 urine samples was carried out using HiChrome UTI agar (M1353R; HiMedia Laboratories Pvt. Ltd.) media following the pour plate method followed by the serial dilution technique. Each of the collected urine samples were diluted as per serial dilution procedure and from each dilution 1ml was directly added to the respective UTI agar media and poured onto the plates, and incubated at 37 ± 2°C for 24 hours. After incubation the appeared bacterial colonies were subjected to sub-culturing by streaking to obtain the pure colony.

### Characterization and identification of the bacteria

The isolated bacterial strains were characterized by looking into the morphological, microscopic and biochemical characteristics. The colony morphology was studied by naked eye observation of the colonies appeared on petri-dishes. The parameters taken were colour of the colony, solubility on water, opacity, elevation, texture and consistency etc. The isolated bacterial strains were subjected to microscopic observation (Olympus CX21i microscope) following the Gram’s staining technique. The isolated bacterial strains were further identified by performing different biochemical tests which includes IMViC tests (Indole, Methyl red, Voge’s Proskauer, Citrate), Oxidase test, TSI agar test (Triple sugar iron), test for H_2_S production and Catalase test. The biochemical tests were performed by following the standard biochemical test protocol (10). The reagents used for these tests were procured from HiMedia Laboratories Pvt. Ltd.

### Antibiotic sensitivity test (AST)

The antibiotic sensitivity test was carried out by following the Kirby-Bauer Disc diffusion method (11). The antibiotics used for this study were Ciprofloxacin, Levofloxacin, Ofloxacin, Ceftriaxone and Piperacillin (HiMedia Laboratories Pvt. Ltd.). The overnight grown bacterial cultures were spreaded onto the nutrient agar plate and antibiotic discs were placed over the media. The plates were allowed to incubate at 37°C for 24 hours. After incubation, the zones of inhibition were measured and compared with the Kirby-Bauer interpretative chart and inferred as resistant, sensitive or intermediately sensitive.

### Assessment of the activity of plant extracts against the UTI pathogens

#### Collection of plant materials

A total of 10 plants from the family Fabaceae (*Tamarindus indica*, *Trigonella foenum-graecum*, *Clitoria ternatea*, *Dalbergia sissoo*), Lamiaceae (*Ocimum scantum*, *Ocimum basilicum*, and *Mentha pipertia)* and Zingiberaceae (*Zingiber officinale*, *Curcuma longa*, *Canna indica*) were collected from different parts of Odisha considering their ethno-medicinal importance.

#### Isolation of crude extracts

Leaves of the plants were washed with distilled water, dried in shade, grounded using mortar and pestle and stored in airtight container at room temperature in dark until used. The powder samples were separately subjected to aqueous extraction. Aqueous extracts were prepared by dissolving 20 grams of powdered leaves in 50 ml of double distilled water such that the level of the solvent was above that of the plant materials. The macerated mixtures were then left on the shaker for 72 hours at room temperature. The extracts were filtered out from the macerated mixture by using Whatman grade 1 filter papers. The aqueous extracts were concentrated using a freeze drier and stored at 4°C for further use.

#### Determining the activity of crude extracts

To evaluate the activity of the leaf extracts on the UTI isolates, ‘well diffusion’ method was followed. The overnight grown bacterial cultures were swabbed onto the solidified nutrient agar plates. Wells were made equidistantly with the help of cork and borer. 30 μl (250μg/ml) of crude extract from each plant was added in respective well on the medium and allowed to stand for 1 hour for proper diffusion and thereafter incubated at 37 °C for 24 hour. All the crude extract samples were studies in triplicates, and the resulting inhibition zones were measured (in mm) and analyzed.

### Dilution susceptibility tests

Dilution susceptibility tests were carried out by broth dilution method (12) to determine the Minimum inhibitory concentration (MIC) and Minimum Lethal Concentration (MLC) values. A series of nutrient-agar broth tubes containing varied concentrations of crude extract (0.062, 0.125, 0.25, 0.5, 1.0, 2.0 mg ml^−1^) were prepared and inoculated with 100 μl inoculums containing 1×10^6^ CFU ml^−1^. Plates were incubated at 37°C for 24 h with one solvent control. The lowest concentration of the crude extract resulting in no growth after 18 to 24 hours of incubation was considered as the MIC. MLC values were also estimated for each crude extract. The lowest concentration of crude extract from which the microorganisms do not recover and grow when transferred to fresh medium was considered as the MLC. The MLC was ascertained if the plates showing no growth are sub-cultured into fresh medium lacking crude extract.

### Statistical analysis

Binary probit model and descriptive statistics were used to analyze the data of 602 clinically suspected UTI patients within the age groups ‘01-75 years’ to determine the UTI prevalence over age groups and gender using the STATA (Version 13) statistical package.

## RESULTS

### Isolation and characterization of the of bacterial strains

The primary UTI agar culture plates were observed with appearance of different types of bacterial colonies. The colonies were of different colours, sizes and textures (Figure 1; Table 1). The characterization of the isolated bacterial strains includes the colony morphology study, microscopy study and biochemical characterization.

**Figure 1.**
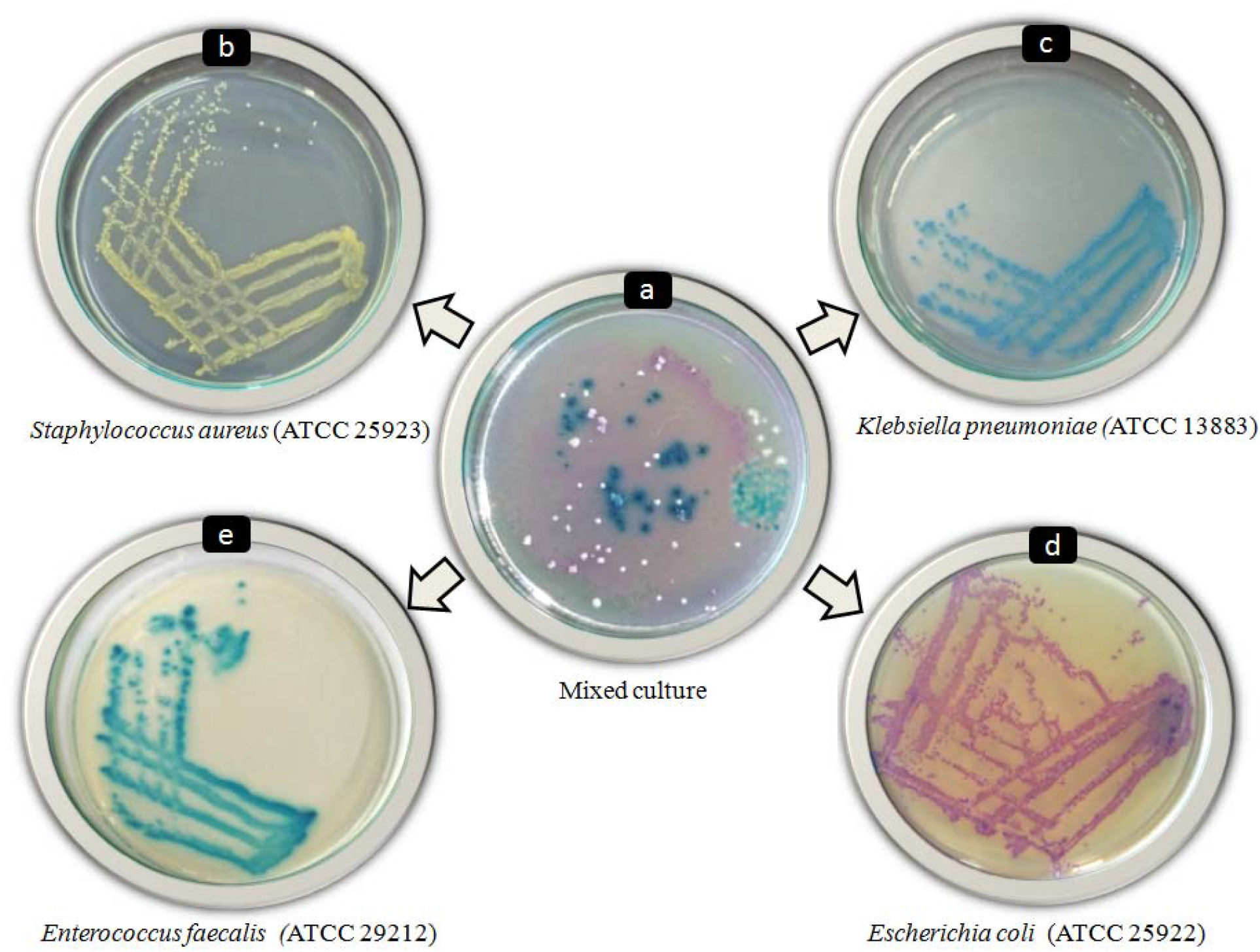
Depiction of the mixed culture of bacterial strains and primarily identified bacterial strains on petri plates. a) Primary culture plate (Mixed culture), b) CSAS-I (Golden yellow: identified as *Staphylococcus aureus*), c) CSAS-F (Deep blue: identified as *Klebsiella pneumoniae*), d) CSAS-G (Pink-purple: identified as *Escherichia coli*), e) CSAS-A (Bluish green: identified as *Enterococcus faecalis*)

**Table 1.** Depiction of colony morphology of the isolated bacterial pathogens causing UTI

| Code | Chromogenesis | Water soluble | Opacity | Elevation | Texture | Size | Consistency | Form | Margin | Appearance | Development |
| --- | --- | --- | --- | --- | --- | --- | --- | --- | --- | --- | --- |
| CSAS-A | Greenish blue | yes | Translucent | Flat | Rough | Large | Mucus | Circular | Entire | Shiny | Dev. Inside |
| CSAS-G | Pink-purple | yes | Opaque | Raised | Smooth | Large | Mucus | Circular | Entire | Dull | On surface |
| CSAS-F | Blue | No | Translucent | Raised | Smooth | Small | Stick to loop | Circular | curled | Shiny | On surface |
| CSAS-I | Golden yellow | No | Opaque | Umbonated | Rough | Puntiform | Stick to loop | Circular | Undulate | Shiny | On surface |

#### Colony morphology study

Four different types of colonies were observed in the primary culture plates and were subjected for sub-culturing. The colonies were coded as CSAS-A, CSAS-G, CSAS-F and CSAS-I. The results of streak plates on UTI agar were depicted in Figure 1. The colony of CSAS-A was greenish blue, translucent, flat with rough texture. CSAS-G showed pink-purple, opaque, raised and smooth colonies. The colony of CSAS-F was deep blue, translucent, raised and smooth where as the colony of CSAS-I was golden yellow, translucent, umbonated with rough surface. The characteristic features of observed colonies are depicted in Table 1. Following the manufacturer’s instruction with references to chromogenesis of colonies on HiChrome UTI agar, CSAS-G was primarily identified as *E. coli* (ATCC 25922). CSAS-F was identified as a strain of *K. pneumoniae* (ATCC 13883). CSAS-I was identified as a strain of *S. aureus* (ATCC 25923) and CSAS-A was identified as strain of *E. faecalis* (ATCC 29212).

#### Microscopic study

The bacteria isolated from urine samples were visualized under compound microscope and the obtained results are represented in Figure 2 and Table 2. The CSAS-A and CSAS-I were observed as Gram’s positive cocci bacterial strains. In contrast to this, the other two bacterial strains namely CSAS-G and CSAS-F were Gram’s negative Bacilli. CSAS-G was hyphen like small rod as compared to that of CSAS-F which was thicker than CSAS-G.

**Figure 2.**
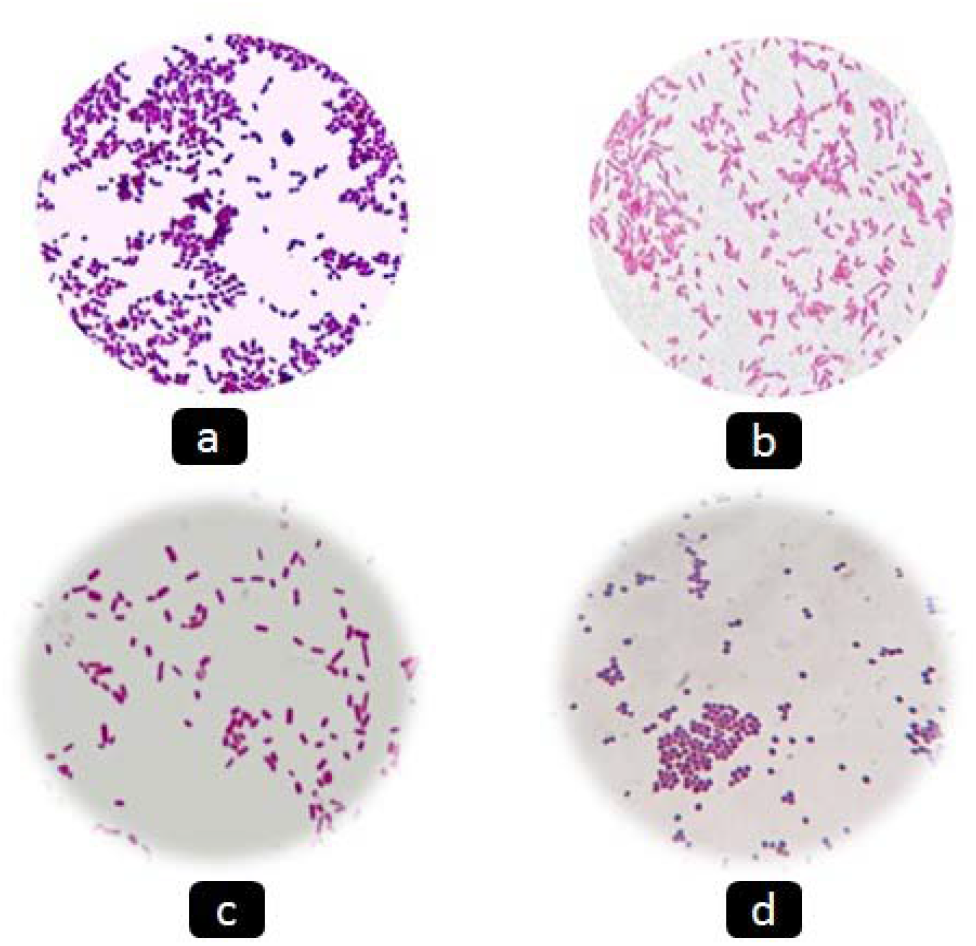
Gram’s stained images of the isolated bacterial strains under 1000x magnification. a) CSAS-A (identified as*E. faecalis*), b) CSAS-G (*E. coli*), c) CSAS-F (*K. pneumoneae*), d) CSAS-I (*S. aureus*)

**Table 2.**
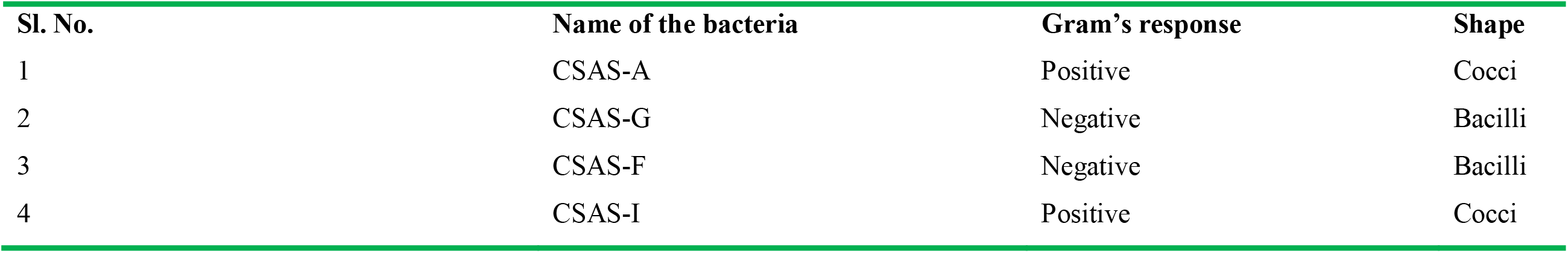
Results of Gram’s staining carried out to identify the bacterial strains

#### Biochemical characterization

The biochemical tests were carried out for confirming the bacterial pathogens preliminarily identified through colony morphology using HiChrome UTI agar (differential media). The results were interpreted by the reaction taken place in the form of change in colour observed by naked eye and are depicted in Figure 3 and Table 3. In the indole test, the strain CSAS-G was showing the formation of cherry red colour ring at interface of medium which indicates the production of indole from tryptophan indicating the positive result. The other three bacterial strains namely CSAS-A, CSAS-F and CSAS-I were giving negative result. The methyl red test identified CSAS-G and CSAS-I as good fermenters of glucose where as CSAS-A and CSAS-F were detected as glucose non-fermenters. CSAS-A, CSAS-F and CSAS-I were positive for VP test by producing pink colour, but CSAS-G was negative. In case of citrate test, CSAS-F and CSAS-I were producing the blue colour which referred to the positive result. All the test bacterial strains were negative for oxidase test. In the triple sugar iron (TSI) test, all the four bacterial strains were detected as fermenters of glucose, lactose and sucrose by producing yellow colour butt and slant which indicates acid production. The isolated bacterial strains were positive for H_2_S production which was evidenced from the production of black colour on TSI agar. All the identified strains were observed to be Catalase positive except CSAS-A.

**Figure 3.**
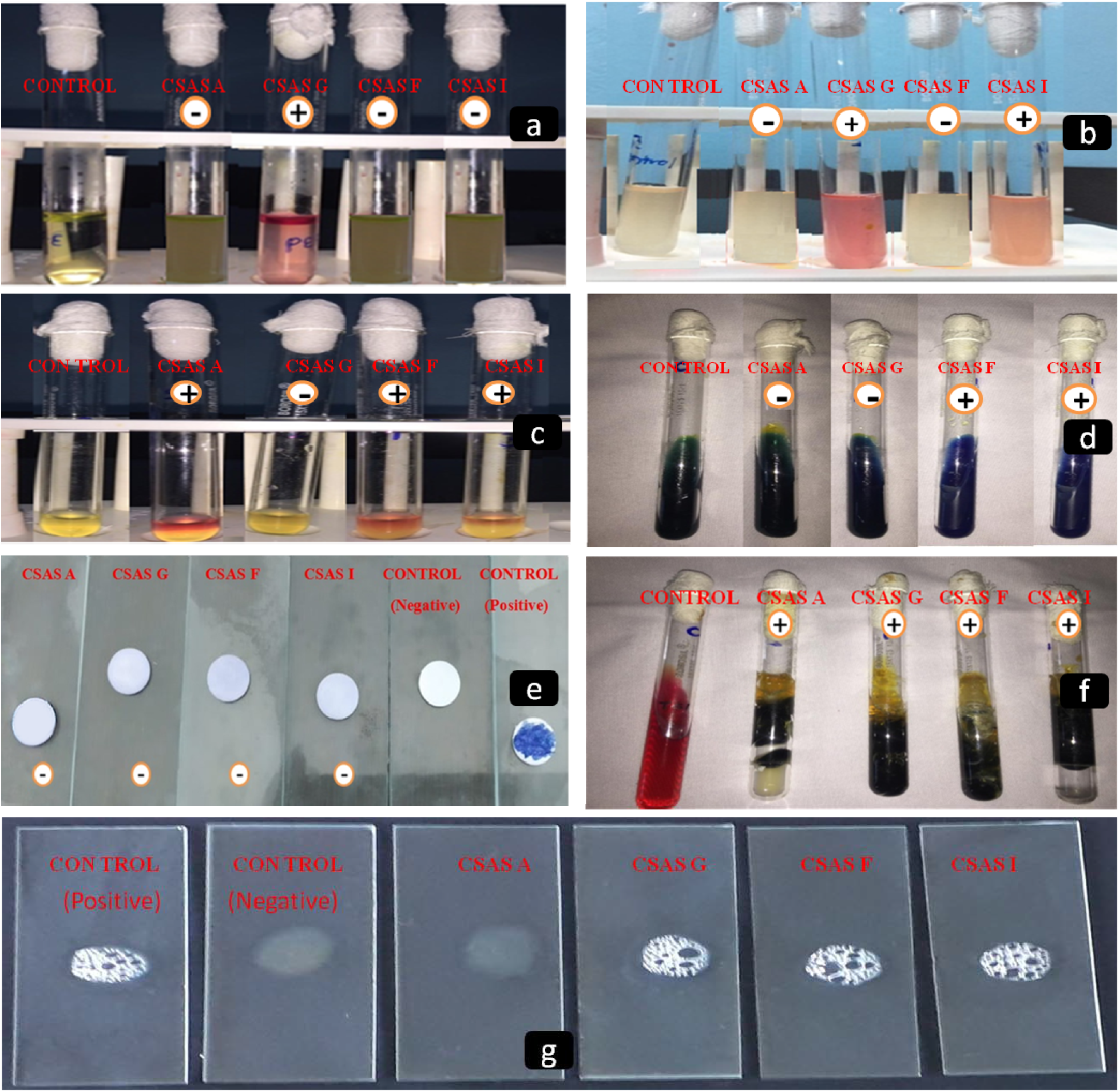
Results of biochemical tests. a) Results of Indole test; positive result is indicated by formation of cherry red colour ring, b) Results of Methyl Red test; positive result is indicated by formation of red colour, c) Result of Voge’s Proskauer test; positive result is indicated by formation of pink colour, d) Result of Citrate Test; positive result is indicated by formation of blue colour, e) Results of oxidase test; positive result is indicated by formation of dark blue colour, f) Results of TSI test; positive result is indicated by formation of yellow colour, g) Resuts of Catalase test; positive result is indicated by formation of bubbles

**Table 3.** Results of biochemical tests carried out to identify the bacterial strains

| Sl. No. | Colony name | Indole | Methyl red | Voge's Proskauer | Citrate | Oxidase | Catalase | TSI | Identified as |
| --- | --- | --- | --- | --- | --- | --- | --- | --- | --- |
| 1 | CSAS-A | - | - | + | - | - | - | G+L+S+ | *E. faecalis* |
| 2 | CSAS-G | + | + | - | - | - | + | G+L+S+ | *E. coli* |
| 3 | CSAS-F | - | - | + | + | - | + | G+L+S+ | *K. pneumoneae* |
| 4 | CSAS-I | - | + | + | + | - | + | G+L+S+ | *S. aureus* |

### Prevalence of pathogens causing UTI among different age groups and gender

A total of 602 urine samples within the age of 01-75 years from clinically suspected patients were analyzed for UTI (Table 4). Among them, 440 (73.09%) samples were found to be culture positive showing significant bacteriuria and the remaining 162 (26.91%) samples were either non-significant bacteriuria or had sterile urine. From the total 440 patients, 286 (65%) female and 154 (35%) male showed culture positive significant bacteriuria. Female gender was a significant risk factor for acquiring UTI compared to male except the female in the age group of 46-75 (Table 5). The prevalence of UTI was highest within the age group of 16-30 years (49%), followed by 31-45 years (28%), among the female patients. Whereas majority of the isolates (48%) were obtained from male patients aged 61-75 years (Table 5). Table 6 illustrates the overall frequency of prevalent pathogens. Out of the 440 significant isolates, Gram-negative accounted for 294 (67%), while Gram-positive accounted for 146 (33%). *E. coli* (60%) was the most pre-dominant isolate causing UTI, followed by *S. aureus* (25%)*, E. faecalis* (8%), and *K. pneumonia* (7%).

**Table 4.**
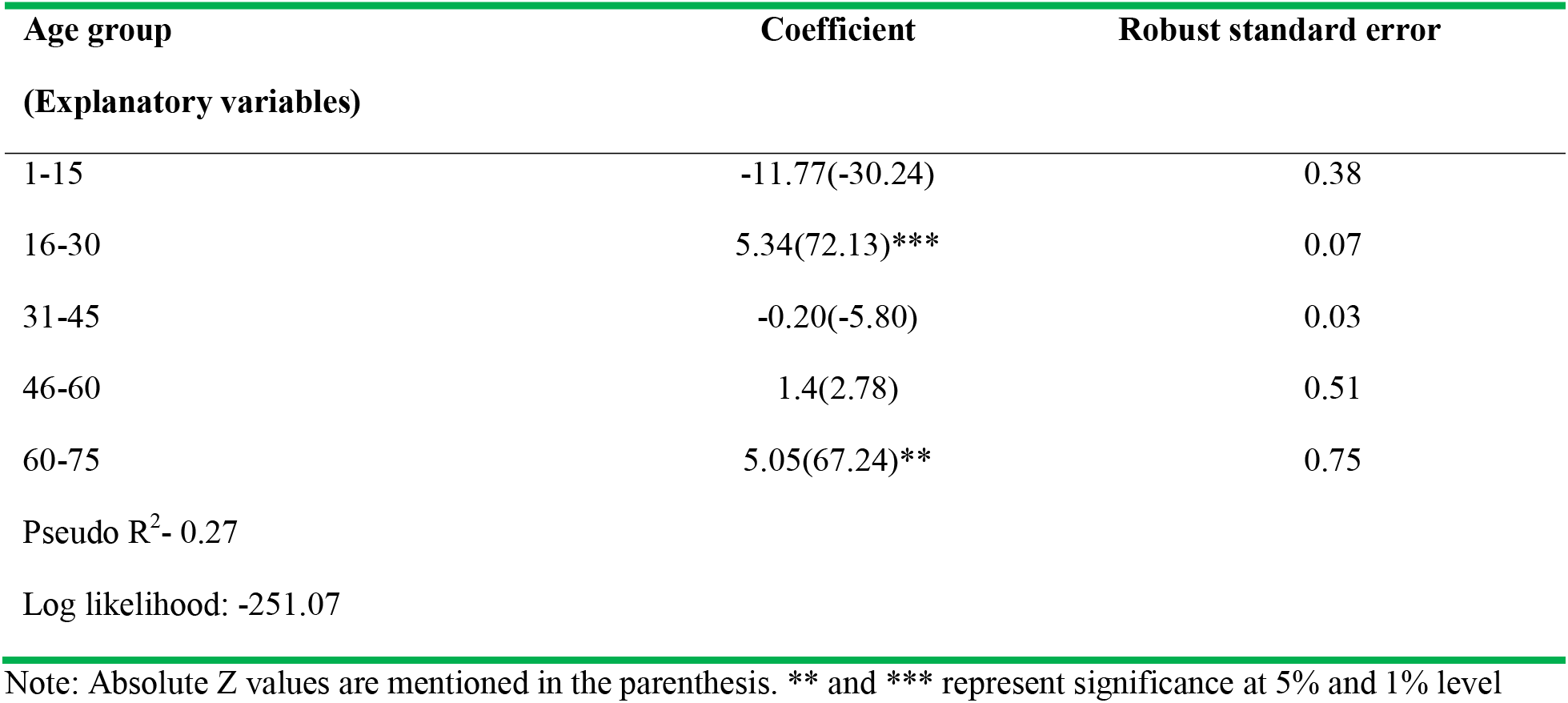
Exposure of different age group of people to urinary tract infection

**Table 5.** Distribution pattern of UTI patients across different age groups and gender

| Age group<br>(in years) | Females<br>(N=286) | Males<br>(N=154) | Mean± SD |
| --- | --- | --- | --- |
| 1-15 | 15(5%) | 8(5%) | 10±0.21 |
| 16-30 | 138(49%) | 22(14%) | 21±0.38 |
| 31-45 | 80(28%) | 25(14%) | 38±0.32 |
| 46-60 | 19(9%) | 31(19%) | 54±0.48 |
| 61-75 | 27(9%) | 74(48%) | 69±0.39 |

Note: Values in the parenthesis represent the percentage.

**Table 6.** Prevalence of uropathogens among males and females

| Uropathogens | Male, n (%) | Female, n (%) | Total, n (%) |
| --- | --- | --- | --- |
| *E. coli* | 85(56%) | 179(62%) | 264 (60%) |
| *S. aureus* | 28(18%) | 82(29%) | 110 (25%) |
| *K. pneumonia* | 13(8%) | 17(6%) | 30 (7%) |
| *E. faecalis* | 28 (18%) | 8(3%) | 36(8%) |
| Total | 154 (100%) | 286 (100%) | 440 (100%) |

### Antibiotic susceptibility test

The antibiotic susceptibility test towards the prescribed antibiotics Ciprofloxacin, Levofloxacin, Ofloxacin, Ceftriaxone and Piperacillin were depicted in Figure 4 and Figure 5. The test plates were observed with well marked zone of inhibition towards Fluoroquinolone group of antibiotics (Ofloxacin, Levofloxacin, and Ciprofloxacin). But, the zones of inhibition were less in case of Cephalosporins (Ceftriaxone) and β-lactam antibiotic (Piperacillin). The zones of inhibition were measured and compared with the Kirby-Bauer chart and inferred as Resistant (R), Sensitive (S) or Intermediately Sensitive (I). The results are depicted in Table 7.

**Figure 4.**
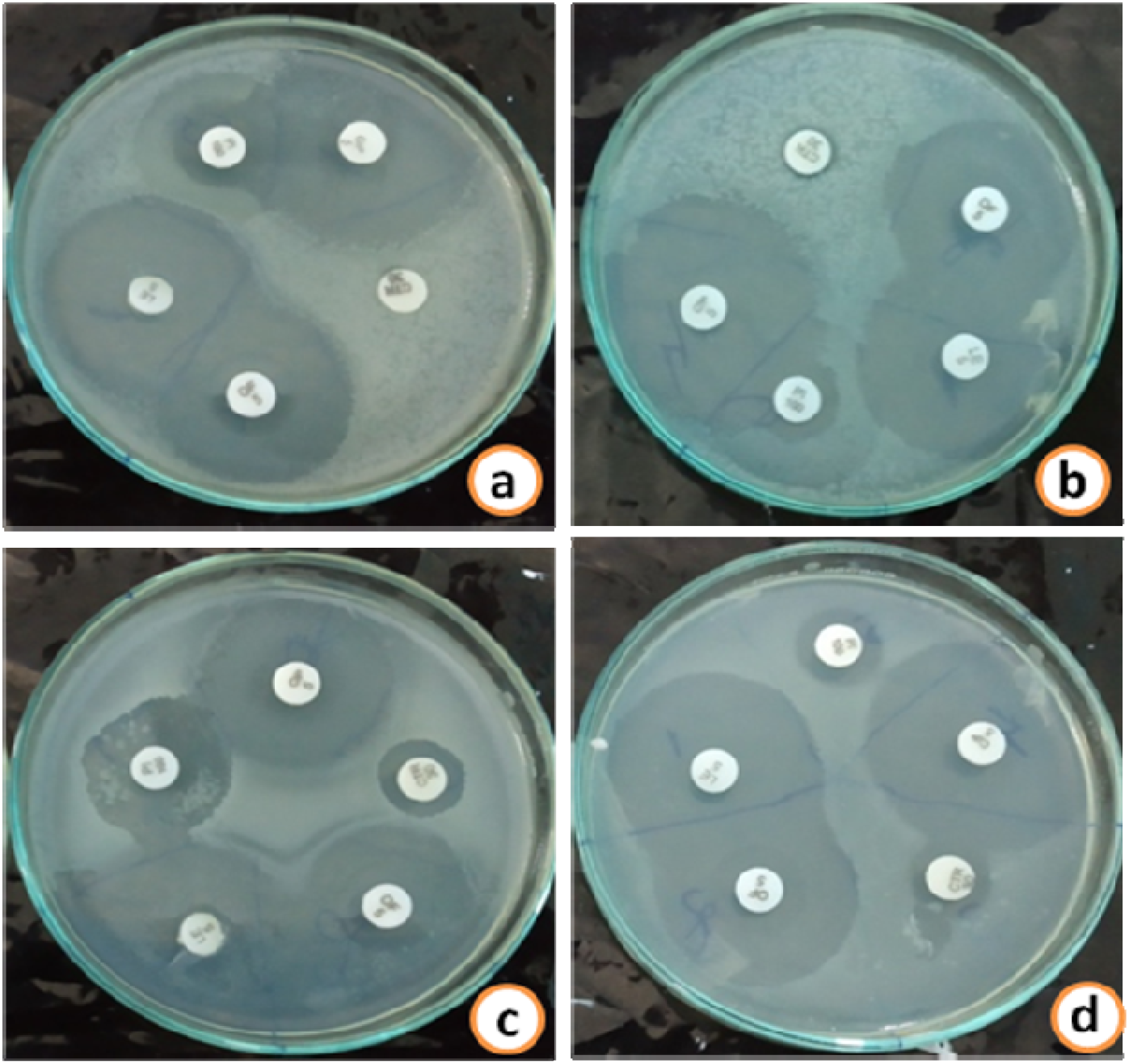
Antibiotic susceptibility test sowing sensitivity of UTI causing pathogens to different antibiotics. CSAS-A (a), CSAS-G (b), CSAS-F (c), and CSAS-I (d); Antibiotics used are Ciprofloxacin (10 μg), Levofloxacin (5 μg), Ofoxacin (5 μg), Ceftriaxone (30 μg), and Piperacillin (100 μg)

**Figure 5.**
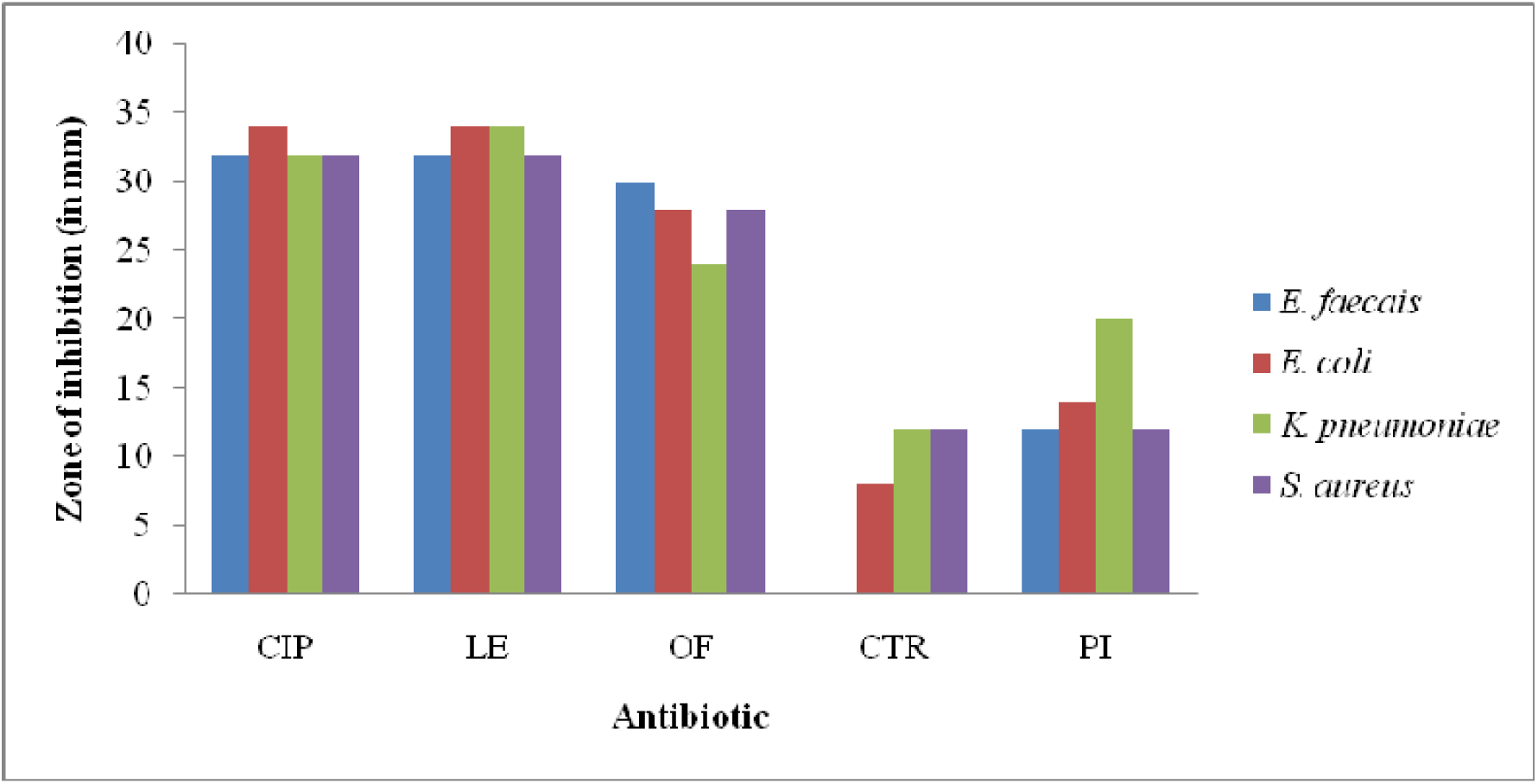
Depiction of the zone of inhibition shown by different antibiotics against the UTI pathogens.

**Table 7.** Antibiotic susceptibility test showing the zone of inhibition (in mm)

| Code No | Identified bacterial strain | CIP (10µg) |  | LE (5µg) |  | OF (5µg) |  | CTR (30µg) |  | PI (100 µg) |  |
| --- | --- | --- | --- | --- | --- | --- | --- | --- | --- | --- | --- |
| CSAS-A | *E. faecalis* | 32 | S | 32 | S | 30 | S | 0 | R | 12 | R |
| CSAS-G | *E. coli* | 34 | S | 34 | S | 28 | S | 8 | R | 14 | R |
| CSAS-F | *K. pneumoneae* | 32 | S | 34 | S | 24 | S | 12 | R | 20 | I |
| CSAS-I | *S. aureus* | 32 | S | 32 | S | 28 | S | 12 | R | 12 | R |

Note: R: Resistant, S: Sensitive, I: Intermediately Sensitive, CIP: Ciprofloxacin, LE: Levofloxacin, OF: Ofloxacin, CTR: Ceftriaxone, PI: Piperacillin

### Assessment of the efficacy of aqueous leaf extracts against UTI isolates

The leaf extracts were tested for antibacterial activity on all the four isolated bacterial strains. The test plates treated with leaf extracts of *T. foenum-graecum*, *D. sissoo*, *O. scantum*, *O. basilicum, M. pipertia*, *Z. officinali*, *C. longa* and *C. indica* were observed with well growth of bacterial strains without any zone of inhibition. On contrary, aqueous leaf extracts of *T. indica* showed zone of inhibition about 24.5±1.29 mm, 23.5±0.58 mm, 22.5±0.58 mm and 20.5±0.58 mm against *E. faecalis, E. coli, K. pneumoniae and S. aureus*, respectively (Figure 6; Table 8). Similarly, *C. ternatea* aqueous leaf extracts also showed zone of inhibition of about 19.25±1.26mm, 17.75±0.96mm, 15.5±0.58mm and 16.5±0.58mm against *E. faecalis, E. coli, K. pneumoniae and S. aureus*, respectively (Table 8). The zones of inhibition shown by these extracts were more than the zones of inhibitions shown by some of the tested antibiotics (Figure 6; Table 8).

**Figure 6.**
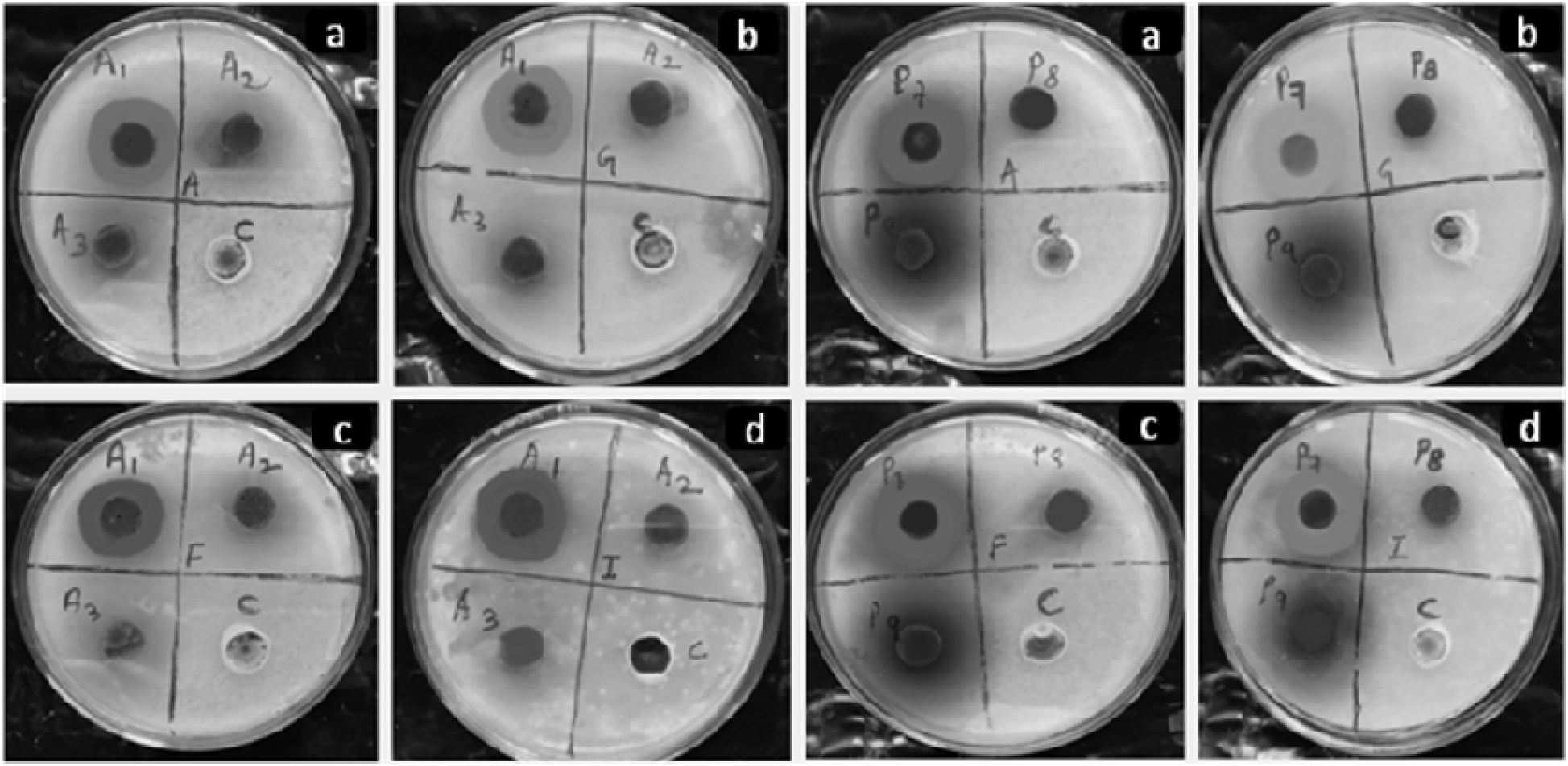
Representative figure showing effect of aqueous leaf extracts on bacterial pathogens *E. faecalis*, *E. coli*, *K. pneumoniae* and *S. aureus* in a, b, c and d respectively. A1, A2 and A3 are the extracts of *T. indica*, *T. foenum-graceum*, and *C. longa*. P7, P8 and P9 are the extracts of *C. ternatea*, *D. sissoo* and *O. scantum*

**Table 8.** Zone of inhibition shown by the aqueous leaf extracts of *Tamarindus indica* and *Clitoria ternatea* tested against the UTI isolates

| Bacterial pathogen | *Tamarindus indica* |  |  | *Clitoria ternatea* |  |  |
| --- | --- | --- | --- | --- | --- | --- |
|  | Inhibition zone at 0.25 mg/ml (in mm)* | MIC (mg/ml) | MLC (mg/ml) | Inhibition zone at 0.25mg/ml (in mm)* | MIC (mg/ml) | MLC (mg/ml) |
| *E. faecalis* | 24.5±1.29 | 0.25 | 0.25 | 19.25±1.26 | 0.125 | 0.25 |
| *E. coli* | 23.5±0.58 | 0.125 | 0.25 | 17.75±0.96 | 0.125 | 0.125 |
| *K. pneumoneae* | 22.5±0.58 | 0.125 | 0.125 | 15.5±0.58 | 0.25 | 0.25 |
| *S. aureus* | 20.5±0.58 | 0.062 | 0.062 | 16.5±0.58 | 0.062 | 0.125 |

\*Mean ± SD (n=4)

### Determination of MIC and MLC values

MIC of *T. indica* leaf extract required was less (0.062 mg/ml) to inhibit the growth of *S. aureus* and was more (0.25 mg/ml) to inhibit the growth of *E. faecalis.* The MIC of *C. ternatea* leaf extract required was also less (0.062 mg/ml) to inhibit the growth of *S. aureus* and was more to inhibit the growth of *K. pneumonia*. In almost all the cases the MIC value was in line with MLC value (Table 8). However, the MLC value was found to be more than the MIC when *T. indica* leaf extract was used for inhibiting the growth of *E. coli*. Likewise, the MLC required was more than the MIC when *C. ternatea* leaf extract was used to inhibit the growth of *E. faecalis*. Except these two cases, the MLC was found to be in concurrent with MIC for inhibiting the growth of UTI pathogens. The MIC and MLC of *T. indica* and *C. ternatea* required for inhibiting the pathogenic bacterial growth are depicted in Table 8.

## DISCUSSION

Prevention of any kinds of diseases particularly the infectious diseases is relies on identification and understanding of the infectious counterpart associated with the diseases. With this outset the study identified four bacterial pathogens causing urinary tract infection among indigenous people of Bolangir district, Odisha, India. On the basis of chromogenesis of colonies on UTI-Agar plates and following the instruction manual of HiChrome UTI differential media (M1353R; HiMedia Laboratories Pvt. Ltd.), the UTI isolates were primarily identified as *E. coli* (ATCC 25922), *E. faecalis* (ATCC 29212), *S. aureus* (ATCC 25923), and *K. pneumoniae (*ATCC 13883). The CSAS-G was identified as *E. coli* as it produced pink colour colonies and this is due to the enzyme ß-D-galactosidase that cleaves the other chromogenic substrate present in the media. This was congruent with the result obtained by Akter et al. (13) while studying the antibiotic sensitivity of pathogens causing UTI. CSAS-F was identified as a strain of *K. pneumoniae* due to the production of blue colonies (14). The strains of the CSAS-A was greenish blue in colour which indicated the presence of *E. faecalis* because one of the chromogenic substrate is cleaved by β-glucosidase possessed by *Enterococci* resulting in formation of blue colonies. *Enterococci* were easily discriminated by their strong turquoise pigmentation and their typical growth on the agar surface (15). CSAS-I was identified to be a strain of *S. aureus* as it produced golden yellow colour colonies.

Microscopic and biochemical characterizations were carried out to ascertain the primarily identified bacterial strains obtained through colony morphology study. The results obtained from the microscopic study indicated the prevalence of both Gram positive as well as Gram negative bacterial strains, with coccus and bacillus forms, in the tested urine samples isolated from patients suffering from UTIs. Mollick et al. (16) and Sewify et al. (17) also reported the abundance of both Gram positive and Gram negative bacterial pathogens (with coccus and bacillus forms) in patients with urinary tract infection. This indicated that the Gram positive as well as Gram negative bacteria are equally responsible for UTI and the immune system might not be discriminating and preventing either of them from the infection process. The biochemical tests results were interpreted by the reaction taken place in the form of change in colour (Figure 3). In the indole test, the strain CSAS-G was showing the formation of cherry red colour ring at interface of medium which indicates the production of indole from tryptophan indicating the positive result (18). The other three bacterial strains namely CSAS-A, CSAS-F and CSAS-I were unable to produce indole from tryptophan giving negative result. As methyl red test is used to detect the fermentation of glucose which indicated by production of red colour (19), here CSAS-G and CSAS-I were detected as good fermenters of glucose where as CSAS-A and CSAS-F were glucose non-fermenters. CSAS-A, CSAS-F and CSAS-I were positive for VP test by producing pink colour, but CSAS-G was negative. In case of citrate test, CSAS-F and CSAS-I were producing the blue colour which referred to the positive result and utilization of citrate as the sole source of carbon (20). The oxidase test is used to identify bacteria that produce cytochrome-c oxidase, an enzyme of the bacterial electron transport chain. When present, the cytochrome c oxidase oxidizes the reagent (tetramethyl-p phenylenediamine) to (indophenols) purple color end product. When the enzyme is not present, the reagent remains reduced and is colorless. The oxidase negative result just means that these organisms do not have the cytochrome coxidase that oxidizes the test reagent. In the present study all the test bacterial strains were negative for oxidase test i.e. these organisms might be respiring using oxidases other than the cytochrome c oxidase in the electron transport process. Further, the TSI identified CSAS-A, CSAS-G, CSAS-F and CSAS-I as fermenters of glucose, lactose and sucrose, which is indicated by production of yellow colour butt and slant which pointed out towards acid production (21). Catalase test usually helps to distinguish between Streptococcus and Staphylococcus. In the present study, all the identified strains were observed to be catalase positive except CSAS-A. Summing up, the biochemical tests inferred that the CSAS-A was IMViC − −+−, Oxidase −ve, TSI +ve, Catalase −ve; which indicates that the strain might be from *E. faecalis* (22). CSAS-G was IMViC ++ − −, Oxidase −ve, TSI +ve, Catalase +ve, which indicates it as a strain of *E. coli* (22). CSAS-F was IMViC − − ++, Oxidase −ve, TSI +ve, Catalase +ve, which indicates it as a strain of *Klebsiella* sp. (23). CSAS-I was IMViC − + ++, Oxidase −ve, TSI +ve, Catalase +ve, which indicates it as a strain of *S. aureus* (23). The identification of the bacterial strains by the colony morphology on HiChrome UTI agar differential media and biochemical based identification were in agreement with each other inferring the same results. After all, the study identified four bacterial isolates as possible causative agents of UTI cases observed in the Bolangir districts of Odisha, India. These organisms were *E. coli*, *K. pneumoniae*, *E. faecalis* and *S. aureus*. These uropathogens are also the common urinary tract infection causing agents as reported earlier (24, 25). Along with this, some studies also identified some other bacterial pathogens such as *Pseudomonas* sp., *S. epidermidis*, and S*treptococcus* sp. in the urine samples of patients with urinary tract infection (23). The differences in the distribution pattern of these UTI causing pathogens might be due to the differences in the environmental conditions and host specificity.

From the 602 urine samples collected from UTI patients, 440 (73.09%) yielded significantly culture positive pathogens. The culture positive rate in the present study was more than the rates obtained from the earlier studies carried out in rural community of Odisha (34.5%) (2), rural community of Nigeria (39.7%) (26), Jaipur, India (17.19%)(27) and Aligarh, India (10.86%) (28). The finding showed that females (65%) had higher prevalence of UTI in comparison to males (35%) in agreement with earlier studies (26, 27, 2). Close proximity of the female urethral meatus to anus, short urethra, and sexual intercourse have been reported as factors influencing the higher prevalence of UTIs in women (29). The age group analysis showed that young female patients in the age range of 16-30 years had highest prevalence rate (49%) of UTI among the females. This result is in agreement with previous studies (28, 27, 2). Among sexually active young women the incidence of symptomatic UTI is high, and the risk is strongly associated with recent sexual intercourse, recent use of diaphragm with spermicide, and a history of recurrent UTIs (30). Elderly males (61-75 years) had higher incidence of UTIs among the males (48%) as compared to elderly females (9%) and is in line with the earlier reports (27, 2). The increased incidence of UTI among males might be due to prostate enlargement and neurogenic bladder in elderly male (31). In this study the prevalence of Gram negative bacterial pathogens was more (67%) compared to the Gram positive pathogens (33%). Among all these pathogens *E. coli* was more prevalent and accounted for 60% of all culture-positive isolates, followed by *S. aureus* (25%)*, E. faecalis* (8%), and *K. pneumonia* (7%). The proportion of *E. coli* species isolated was similar to those described in previous studies (32, 27). While the proportion of *S. aureus* (25%)*, E. faecalis* (8%), and *K*. *pneumonia* (7%) contradicts the findings of earlier studies (32, 27, 2). Geographical location of isolated pathogens and/or climate change might be the reason(s) for this kind of differences with respect to the prevalence and distribution of these pathogens.

The antibiotic sensitivity testing is usually done to determine the effectiveness of antimicrobials/antibiotics in inhibiting the growth of the pathogens causing a specific infection. The results from this test will help a healthcare practitioner determine which drugs are likely to be most effective in treating a person’s infection. The present study assesses the effectiveness of the presently prescribed antibiotics such as Ciprofloxacin, Levofloxacin, Ofloxacin, Ceftriaxone and Piperacillin. Among these, the Fluoroquinolone group of antibiotics (Ofloxacin, Levofloxacin, and Ciprofloxacin) represented highest zone of inhibitions towards all the four bacterial pathogens tested as compared to Cephalosporins (Ceftriaxone) and β-lactam antibiotic (Piperacillin), which are mostly prescribed for UTI now-a-days (Figure 5; Table 7) (33). It seems that the UTI isolates (except *Klebsiella* sp.) were more sensitive towards Ciprofloxacin, Levofloxacin and Ofloxacin, and were resistant towards Ceftriaxone and Piperacillin, and intermediately sensitive to Piperacillin (Figure 5). This finding was also supported by Ganapathy et al. (34). This substantiates the need of alternative therapeutics to counteract the effects caused by UTI isolates.

It is the need of the hour to develop new antimicrobials and therapeutic agents having high effectiveness with no side effects, easy availability and less expensive (23). Recently, the exploitation of wild plants for medicinal purposes has gained ample acceptance worldwide. Therefore, in this study we have attempted to study the effectiveness of 10 plants with medicinal properties against the isolated UTI isolates. The leaf extracts were tested for antibacterial activity on all the four isolated bacterial strains. Out of the ten extracts evaluated, only two *viz.* aqueous leaf extracts of *T. indica* and *C. ternatea* showed zone of inhibition against the pathogens. The effect of *T. indica* on growth of UTI causing pathogens has also been studied earlier (35) but was tested only on pathogens causing UTI in women patients. Again, they had used acetonic, methanolic and chloroform leaf extracts to evaluate their antibacterial activity. They found that the acetone extract of *T. indica* leaves showed highest antibacterial activity against *E. coli* (22.5 mm). Methanol extract showed highest activity against *P. aeruginosa* (21.4 mm) and chloroform extract showed highest activity against *Klebsiella* sp. (6.2 mm). In the present study attempt had been made to study the effect of aqueous leaf extracts of *T. indica* on the UTI causing pathogens and zone of inhibition of about 24.5mm, 23.5mm, 22.5mm and 20.5mm were found against *E. faecalis, E. coli, K. pneumoniae and S. aureus*, respectively (Figure 6; Table 8). Further, the occurrence of antibacterial activity by aqueous extracts provides the scientific basis for using this plant in the treatment of bacterial diseases. Similarly, *C. ternatea* aqueous leaf extracts also showed zones of inhibition of about 19.25, 17.75mm, 15.5mm and 16.5mm against *E. faecalis, E. coli, K. pneumoniae and S. aureus*, respectively (Table 8). The zones of inhibition shown by both of these aqueous leaf extracts were more than the zones of inhibitions shown by some of the tested antibiotics such as Ceftriaxone and Piperacillin (Figure 6; Table 8). This might be due to antibiotic resistance shown by the pathogens against the frequently used antibiotics but not against the aqueous leaf extracts. This indicates that there must be having one or more bio-active compound(s) in the leaf of *T. indica* and *C. ternatea* that makes these pathogens sensitive. Various similar kinds of studies were also performed to screen the activity of crude extracts of *C. ternatea* against microbial pathogens and antimicrobial property of this pant has been ascertained (36).

The MIC is the lowest concentration of a drug that prevents growth of a particular pathogen. Some idea of the effectiveness of a chemotherapeutic agent against a pathogen can be obtained from the minimal inhibitory concentration (MIC). The lowest concentration of the chemotherapeutic agent or antibiotic resulting in no growth after 18 to 24 hours of incubation is the MIC. The minimal lethal concentration (MLC) is the lowest drug concentration that kills the pathogen. The MLC can be ascertained if the tubes showing no growth are sub-cultured into fresh medium lacking antibiotic or chemotherapeutic agent. The lowest antibiotic concentration from which the microorganisms do not recover and grow when transferred to fresh medium is the MLC. In the present study, the MIC of *T. indica* leaf extract required was low (0.062 mg/ml) to inhibit the growth of *S. aureus* and was high (0.25 mg/ml) to inhibit the growth of *E. faecalis*. The MIC of *C. ternatea* leaf extract required was also less (0.062 mg/ml) to inhibit the growth of *S. aureus* and was more to inhibit the growth of *K. pneumoniae*. In almost all the cases the MIC value was in line with MLC value (Table 8). However, the MLC value was found more than the MIC when *T. indica* leaf extract was used for inhibiting the growth of *E. coli*. Likewise, the MLC required was more than the MIC when *C. ternatea* leaf extract was used to inhibit the growth of *E. faecalis*. Except these two cases, the MLC was found to be in concurrent with MIC for inhibiting the bacterial pathogens causing the UTI (Table 8). The highest MIC required to inhibit the growth of *Staphylococcus aureus* in *Tamarindus indica* leaf extract might be an indication that the bacteria possesses the capability of acquiring antibiotic resistance particularly for the antimicrobial(s) present in *T. indica* while the lowest MIC required for inhibiting the growth of the same bacteria in *C. ternatea* leaf extract indicates the potentiality of the antimicrobial(s) present in *C. ternatea* leaf extract. In another way it can be stated that a particular bacteria may be sensitive to a specific chemotherapeutic agent or plant extract but may not be to another.

With the foregoing discussion in a nutshell it can be stated that the UTIs are a serious health problem throughout the world next to respiratory infection affecting millions of people each day. The study advocates the prevalence of a diverse group of bacterial pathogens in the urine sample of the patients infected with urinary tract infections (UTIs). The bacterial strains identified by phenotypic studies (colony morphology, microscopic study and biochemical characterization) were *E. faecalis, E. coli, K. pneumonia*e and *S. aureus.* This could be further confirmed by the 16s rDNA sequencing. The finding showed that females (65%) had higher prevalence of UTI in comparison with males (35%). The age group analysis showed that young female patients in the range of 16-30 years had highest prevalence rate (49%) of UTI among the females. Elderly males (61-75 years) had a higher incidence of UTI among the males (48%) when compared with the elderly females of the same age group (9%). Further, the prevalence of Gram negative bacterial pathogens was more (67%) compared to the Gram positive pathogens (33%). Among all these pathogens *E. coli* was more prevalent accounted for 60% of all culture-positive isolates followed by *S. aureus* (25%)*, E. faecalis* (8%), and *K. pneumonia* (7%).

The bacterial strains were found to be resistant towards the tested antibiotics like Piperacillin and Ceftriaxone, but found sensitive towards fluoroquinolone group of antibiotics such as Ofloxacin, Levofloxacin, and Ciprofloxacin. Moreover, all these pathogens became sensitive to the aqueous leaf extracts of *T. indica* and *C. ternatea* with clear zones of inhibitions. This substantiates that the leaf of these plants may contains one or more bioactive compound(s) that preventing the growth of these pathogens. The leaves of these plants may be used for herbal drug formulation for treatment of urinary tract infection as these leaves are non-toxic. To predict the exact compound(s) showing efficacy against these pathogens, characterization of the crude extracts and subsequent evaluation of the characterized compounds either in isolation or in combination against these pathogens (*in vitro* as well as *in vivo*) is needed. Further, frequent use of particular antibiotics against specific pathogen(s) should not be practiced rather new antibiotics or natural therapeutics could be explored and be precisely utilized in medical practices to reduce the antibiotic resistance shown by the pathogens.

## ACKNOWLEDGMENTS

The authors acknowledge the Centurion University of Technology and Management, Odisha, India for the laboratory facility provided and thankful to Mr. Pradeep Sarangi (Regional Director, CUTM, Bolangir) for giving motivation to conduct the study.

## ETHICAL APPROVAL

Approval was obtained from the ethics committee of the Centurion University of Technology and Management, Odisha, India.

